# Identifying a TFIID interactome via integrated biochemical and high-throughput proteomic studies

**DOI:** 10.1101/111682

**Authors:** Wei-Li Liu, Lihua Song, Gina Dailey, Anna Piasecka, Robert A. Coleman

## Abstract

The core promoter recognition TFIID complex acts as a central regulator for eukaryotic gene expression. To direct transcription initiation, TFIID binds the core promoter DNA and aids recruitment of the transcription machinery (e.g., RNA polymerase II) to the transcription start site. Many transcription factors target TFIID to control vital cellular processes. Current studies on finding TFIID interactors have predominantly focused on transcription factors. Yet, a comprehensive interactome of mammalian TFIID has not been established. Therefore, this study sought to reveal potential TFIID-nucleated networks by identifying likely co-regulatory factors that bind TFIID. By using intact native human TFIID complexes, we have exploited three independent approaches including a high-throughput Next Generation DNA sequencing coupled with proteomic analysis. Among these methods, we found some overlapping and new candidates in which we further assessed three putative interactors (i.e., Sox2, H2A and EMSY) by co-immunoprecipitation assays. Notably, in addition to known TFIID interactors, we identified a number of novel factors that participate either in co-regulatory pathways or non-transcription related functions of TFIID. Overal, these results indicate that, in addition to transcription initiation, mammalian TFIID may be involved in broader regulatory pathways than previous studies suggested.

## Introduction

Eukaryotic transcription requires highly coordinated interactions between the transcription initiation machinery and a variety of transcription factors to ensure a proper response to various physiological cues [1]. To accurately transcribe a protein-coding gene, a pre-initiation complex (PIC) containing over 80 different polypeptides is formed at specific regions of the promoter DNA [1]. TFIID, a key component within the PIC, is responsible for recognizing and binding specific promoter DNA sequences (i.e. the core promoter elements). Disruption in TFIID’s activities poses a severe threat to proper cellular functions and hastens disease progression [2-5]. Human TFIID (∼1.2 MDa) consists of the TATA-binding protein (TBP) and 14 evolutionarily conserved TBP-associated factors (TAFs). Once TFIID alights on DNA, it directs recruitment of six other basal transcription factors including RNA Polymerase II to initiate transcription. To properly respond to diverse physiological cues and activate select gene expression programs, sequence-specific DNA binding activators stimulate transcription, in part by targeting TFIID and aiding in its recruitment to the promoter [6-14]. TFIID thereby serves as a co-activator and binds many critical activators [15]. In addition to promoter recognition and co-activator functions, TFIID can both write and read a histone code on chromatin to modulate transcription [16-19].

Many cellular processes involving DNA binding factors are co-regulated. These regulatory pathways often cross-talk and form sophisticated networks in cells. Currently, the mammalian TFIID-nucleated network is poorly characterized. Thus far, the 3’ end messenger RNA processing pathway was found to be co-regulated via TFIID’s recruitment of the cleavage-polyadenylation specificity factor (CPSF) complex during transcription initiation assembly [20]. This work indicates a transcription initiation-coupled polyadenylation pathway directed by TFIID. A recent discovery revealed that the lysyl oxidase-like LOXL2 protein interacts with four of the TFIID subunits [21]. Importantly, LOXL2 mediates lysine oxidation of the TAF10 subunit and thereby inhibits TFIID-mediated gene expression involved in pluripotency [21]. Furthermore, a previous comprehensive study on discovering yeast TFIID interactors by multidimentional mass spectrometry documented an array of distinct factors including proteins involved in RNA processing and signal transduction [3]. Their discovery suggests that yeast TFIID may utilize different TAF subunits to associate with different factors participating in various cellular pathways. However, mammalian TFIID’s involvement in other cellular processes remains unclear. Therefore, to advance our understanding of TFIID-mediated eukaryotic transcription, this study aimed to reveal the spectrum of potential TFIID interactors including those non-transcription related factors. One goal was to identify potential co-regulatory bridging factors involved in other cellular pathways, which may regulate TFIID’s activities in promoter recognition and transcription initiation. We anticipate that these intriguing findings will lead to future discoveries of novel transcription initiation-coupled co-regulatory pathways directed by TFIID. It would be interesting to understand how these multiple activities of TFIID are coordinated.

Currently, a number of limitations exist for conventional approaches to reveal novel protein-protein interactions. In particular for large multi-subunit protein complexes, it is difficult to identify their low abundant/cell type-specific interacting factors or proteins exhibiting transient interactions with their targets. Conventional approaches include conducting mass spectrometry after immunoprecipitations, GST pull-down like strategies or conventional protein purification coupled with functional assays. Results from these approaches can vary based on how those large multi-subunit protein complexes (e.g. TFIID) are generated using different chromatography strategies and immunuopurification procedures. Therefore, in addition to conducting immunoaffinity purification of native TFIID followed by MudPIT mass spectrometry, we also developed a unique, unbiased and less labor-intensive proteomic approach by using the intact 14-subunit TFIID complex as bait to identify potential novel TFIID interactors.

Our newly established approach incorporates an *E. coli* Flagella random 12-mer peptide display screening system (Invitrogen) followed by either conventional sequencing or massive parallel Next Generation sequencing (Ion Torrent) prior to proteomic analysis. This high-throughput method has allowed us to detect at least several hundred thousands of peptides binding to TFIID. We further tested several peptides to examine their contact surfaces within TFIID via label transfer assays. Interestingly, the results revealed that some peptides appeared to target distinct locations on TFIID. Next, we utilized these TFIID-interacting peptide sequences to search for putative TFIID interactors via blast searches against a human coding protein database (uniProtKB). The results from our three protesomic appraoches clearly showed some overlapping categories such as chromatin remodeling factors. Remarkably, in addition to well-known TFIID-binding proteins, we have found a number of novel proteins that participate in various cellular processes in which we have classified all of the candidates into different categories. As a mean to strengthen our findings, four candidates (i.e., Sox2, EMSY, and histone H2A) were selected to verify their direct association with TFIID via co-immunoprecipitation experiments. Overall, our work revealed a spectrum of distinct factors that target TFIID. These unknown TFIID interactors could potentially lead to future discoveries of novel co-regulatory pathways linked to transcription initiation driven by TFIID.

## Results

### Identification of novel proteins associated with native human TFIID complex

#### Flagella random peptide display screening/conventional DNA sequencing approach

We sought to identify potential novel human TFIID-interacting factors that could participate either in co-regulatory pathways or non-transcription related functions of TFIID. To this end, we utilized a peptide screening system consisting of a random 12-amino acid peptide library displayed on the bacterial cell surface Flagella (FliTrx random peptide display, Invitrogen). This method allowed us to search for specific peptides interacting with the surface of the large multi-subunit human TFIID complex (Figure 1). In brief, we first generated homogenous native human TFIID complex by carrying out a highly specific immunoprecipitation from fractionated HeLa nuclear extracts using a peptide-elutable monoclonal antibody (mAb) against the TAF4 subunit of TFIID as previously described [10] (Figure 1, left panel). Next, this highly purified TFIID complex was deposited on a tissue culture dish prior to incubation of the primary FliTrx peptide library bacterial culture. After the incubation, unbound bacteria were gently washed away and then the remaining cells were collected by vortexing. This eluted bacteria were grown and re-incubated this pooled culture with TFIID prior to collection of the TFIID-bound cells. To enrich the peptide pool specifically bound to TFIID, we performed another four rounds of panning as described above. Single colonies were obtained from the fifth panning. To reveal the TFIID-interacting peptides, plasmid DNA from each colony was extracted individually and subjected to DNA sequencing.

**Figure 1.**
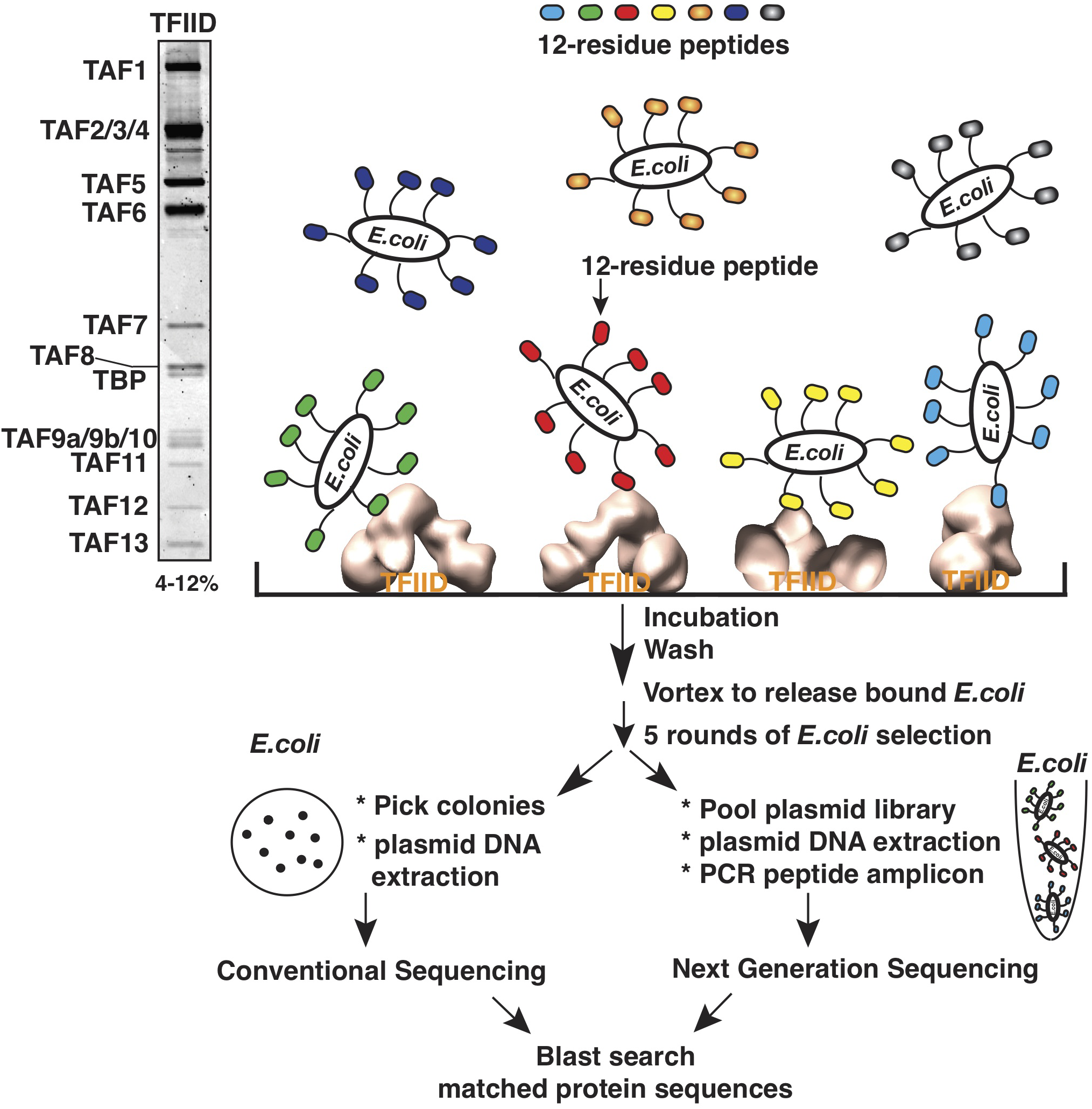
Schematic representation for screening potential TFIID interacting peptides using FliTrx peptide display (Invitrogen). Library of random 12 amino acid peptides, expressed as a thioredoxin fusion on the flagella of *E. coli*, is passed over a petri dish coated with highly purified human TFIID complex isolated from HeLa cells. After 5 rounds of panning, plasmid DNA is purified and subjected to conventional (left) or Next Generation (right) sequencing to determine the identity of TFIID interacting peptides.

Initially we manually sequenced 78 plasmids and obtained 45 in-frame peptides harboring no stop codon. We utilized these in-frame peptide sequences to identify potential TFIID interactors containing these specific peptide sequences by conducting blast searches against a human nuclear protein database (uniProtKB). Interestingly, the results showed that in addition to the TFIID subunits and some known TFIID-binding proteins (e.g. TIF1A, PAX3, RUVBL1) [22-24], a number of novel proteins may associate with TFIID (Table 1). To best illuminate the results, we have classified these proteins into 24 different categories such as Activators/Repressors, RNA Pol II, Mediator, Chromatin remodelers, Histone deacetylases, Histone methylase/demethylase/deimininase complexes, Elongation factors, Splicing factors, RNA capping and cleavage factors, and Pol I/Pol III transcription machinery. Intriguingly, among these 45 peptides, we isolated a rare peptide, designated as D1, composed of 26 amino acids (Supplemental Figure S1). This peptide size was extraordinarily unexpected, given that over 99.5% of the peptide library corresponds to 12 amino acid inserts and was probably the result of a double insertion event during library generation. This D1 peptide has strong homology to a conserved region within the <u>H</u>igh <u>M</u>obility <u>G</u>roup (HMG) box of Sox2 (Figure S1). This rare 26 amino acid D1 peptide isolated in our screen suggests that a strong selection of this sequence was achieved. Thus, this result may identify a novel TFIID interaction region within the Sox2 HMG box domain, consistent with previous studies showing that select HMG domains from other proteins contact TFIID [25-27].

**Table 1.**
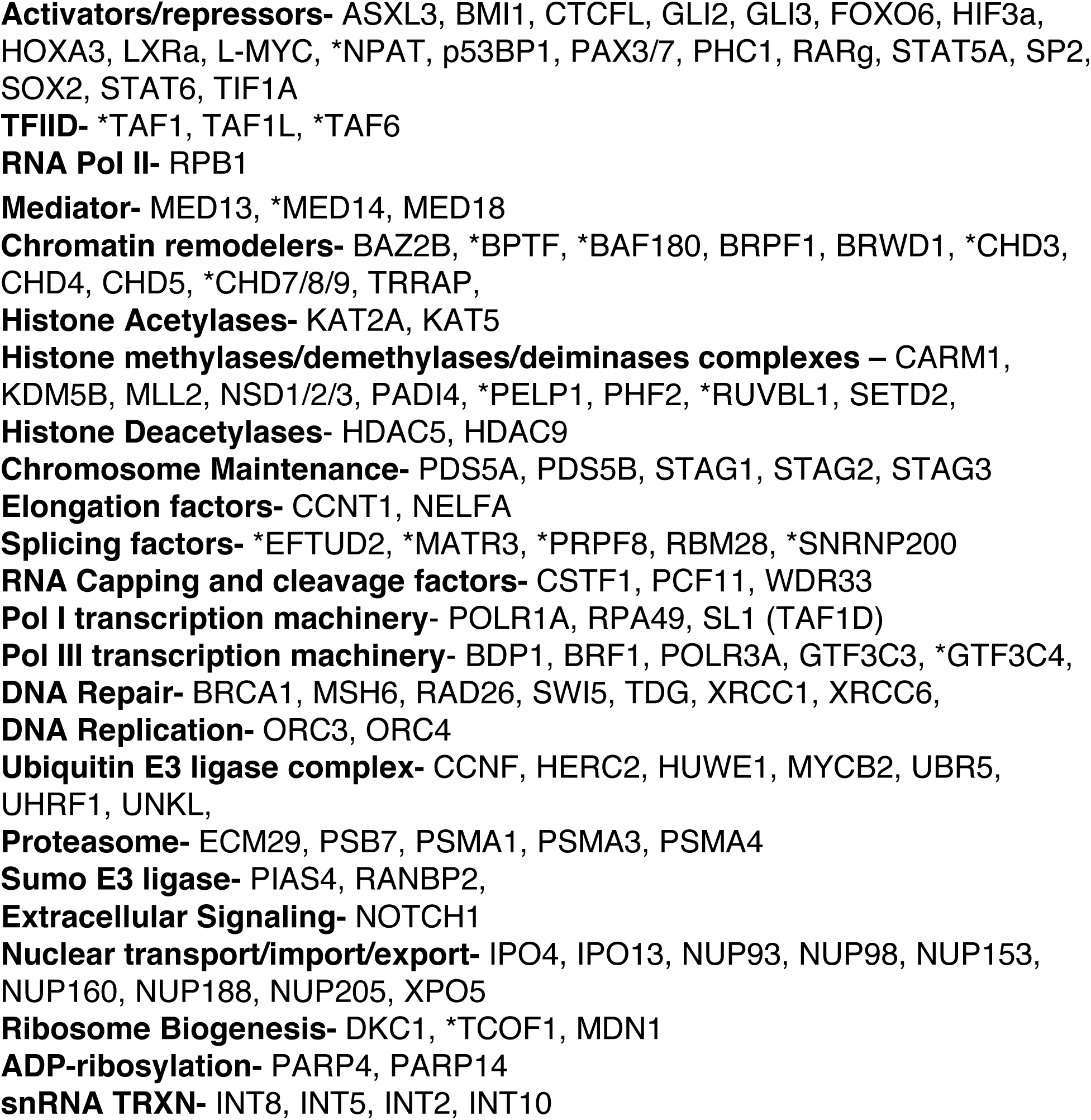
Known/putative TFIID-interacting factors identified based on 45 in-frame peptides. Blast search was conducted following the Flagella random peptide display screening and conventional sequencing. Based on their primary cellular functions, factors analyzed were listed in different categories. Hits also found in MudPIT (Supplemental Table 1 and Table 2) are indicated with a star (*).

#### Immunoaffinity purification of TFIID for MudPIT mass spectrometry

In parallel to the previous approach, we also generated highly purified native human TFIID complex for quantitative MudPIT mass spectrometry [10] to identify novel TFIID interactors. This also allows us to examine the coverage of candidates listed in Table 1. For better evaluation of the results, we performed additional mock immunoprecipitations to detect nonspecific contaminants. In addition, we analyzed two independent TFIID preps to define strong overlapping positive candidates (Table 2). The raw MudPIT analysis is shown in Supplemental Table 1. Table 2 showed that co-precipitates comprised TFIID subunits along with multiple subunits from known TFIID interactors such as the PBAF Chromatin remodeling complex. Intriguingly, we also detected some proteins that were not previously known to interact with TFIID, including factors involved in chromatin remodeling (e.g. PCAF/STAGA), Splicing/RNA processing (e.g. SNRNP200), Ribosome biogenesis (e.g. NOP9), Replication (CTF8), RNA Pol III transcription (e.g. GTF3C4) and Ubiquitination (e.g. SKP1). Next, when comparing results from both methods, we clearly found some overlapping classes that include TFIID, mediator, chromatin remodelers, histone methylation, mRNA processing and RNA Pol III transcription. The factors specifically detected from both methods were highlighted in Table 1. The spectrum of positive candidates (Tables 1 & 2) obtained from both methods indicates that TFIID might associate with more factors involved in a number of distinct regulatory pathways than currently known. In addition, these results suggest that this Flagella random peptide screening/DNA sequencing method is applicable for finding novel factors that interact with multi-subunit complexes (e.g. TFIID in this study).

**Table 2.**
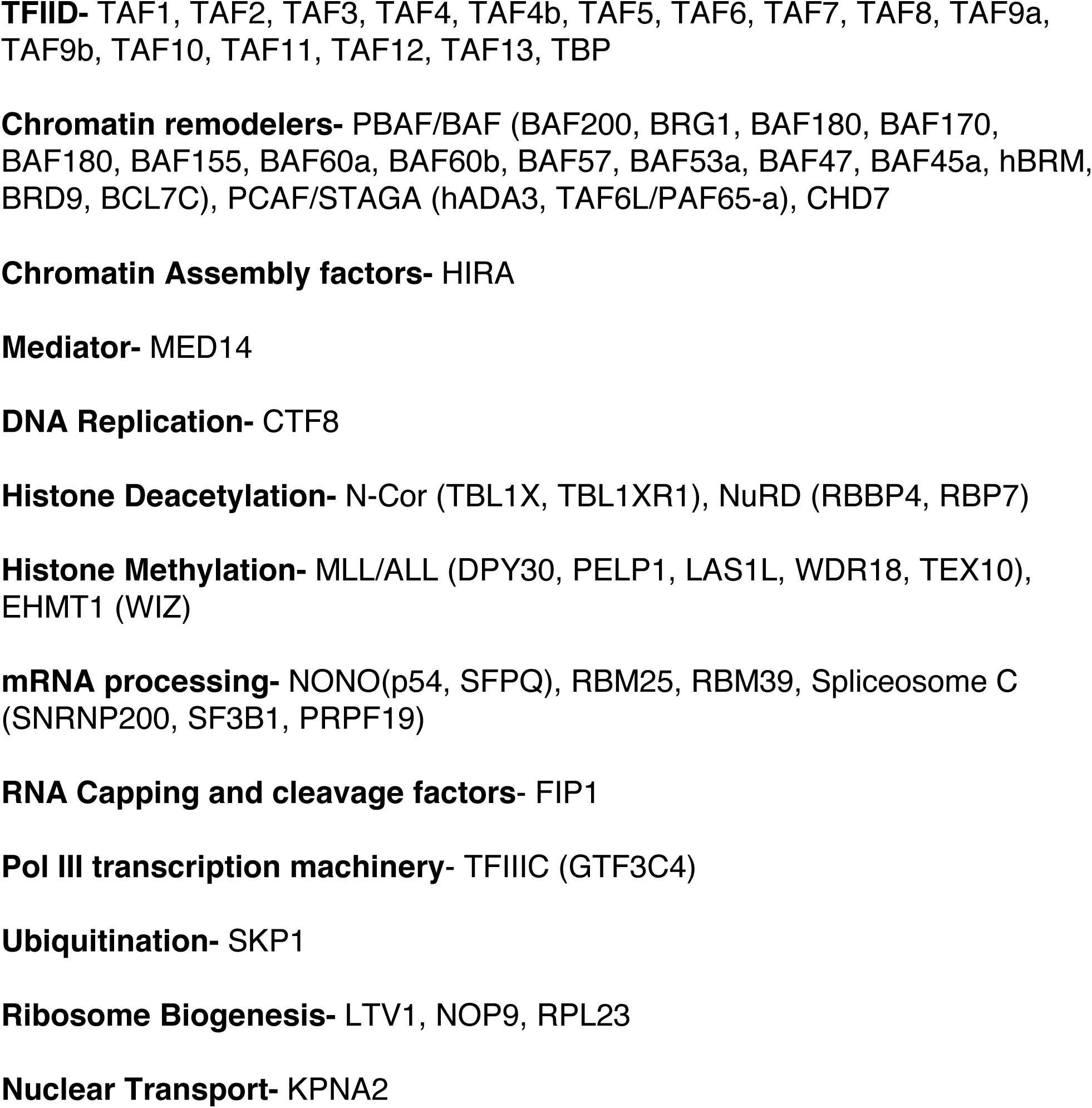
Known/putative TFIID-interacting factors identified based on MudPIT. Blast search was conducted following the affinity immunopurification of native TFIID and MudPIT analysis. Strong overlapping positive hits were identified from the peptides with presence in two independent TFIID preps and filtered with NSAF value above 10^-4^ and 5 times more than control NSAF. Based on their primary cellular functions, factors analyzed were listed in different categories.

#### High-throughput detection of common TFIID-interacting peptide sequences and interactors

Thus far, our results suggest that the Flagella random peptide display screening/sequencing method is feasible for identifying novel protein interactors with large protein complexes. However, manual colony picking/plasmid purification followed by conventional sequencing is labor- and time-consuming, which directly limits the spectrum of proteins identified. Therefore, we exploited an alternative facile approach using Next-Generation DNA sequencing (Ion Torrent, Thermo Fisher). This significantly reduces the high demands of labor and time associated with the conventional approach. In brief, we conducted the same peptide screening experiment (Figure 1). After 5 rounds of enrichment of TFIID-bound bacteria, we directly extracted plasmids from this pooled bacteria followed by PCR amplification. The PCR fragments encoding these peptides were then subjected to Next-Generation sequencing. Importantly, to better validate this approach and help rule out false positive hits, we also performed two additional control experiments including the peptide-library alone (i.e. no input target protein) and a monoclonal antibody against the N-terminus of the largest TFIID subunit TAF1 in which its precise epitope hasn’t been determined yet.

Next-Generation DNA Sequencing yielded large sequence datasets (Figure 2). We assembled these DNA sequences into contigs (a group of two or more identical DNA sequencing reads), followed by translation of individual contig sequences into peptide sequences (Geneious). Peptide sequences containing stop codons and frame shifts were removed from the analysis. We rationalized that a particular peptide sequence was enriched when it had a larger number of sequencing reads than in the non-selected library. Indeed, the total number of contigs assembled from the TFIID and anti-TAF1 mAb selected libraries are significantly higher than the contigs obtained for the non-selected library (Figure 2A). Importantly, this analysis can quantitatively define the number of times a particular peptide appears within the TFIID- or anti-TAF1 mAb-enriched peptide population. Since TFIID is a large protein complex (∼1.2 MDa), we expected that the surface of TFIID could bind more diverse peptides than the anti-TAF1 mAb (∼150 KDa). When we analyzed the top 500 contigs from these three experiments, the highest number of the same-sequence TFIID-interacting peptides is 14 hits, whereas the repeats from the positive control of anti-TAF1 mAb is 235 hits and the negative control with the library alone is only 4 hits (Figure 3A). In addition, approximately 70% of the top 500 TFIID-enriched peptide contigs contain 4 sequencing reads versus 1.2% for the non-selected library (Figure 3B). Collectively these results show that our screening/analysis is specific to the target proteins (i.e. TFIID v.s. anti-TAF1 mAb).

**Figure 2.**
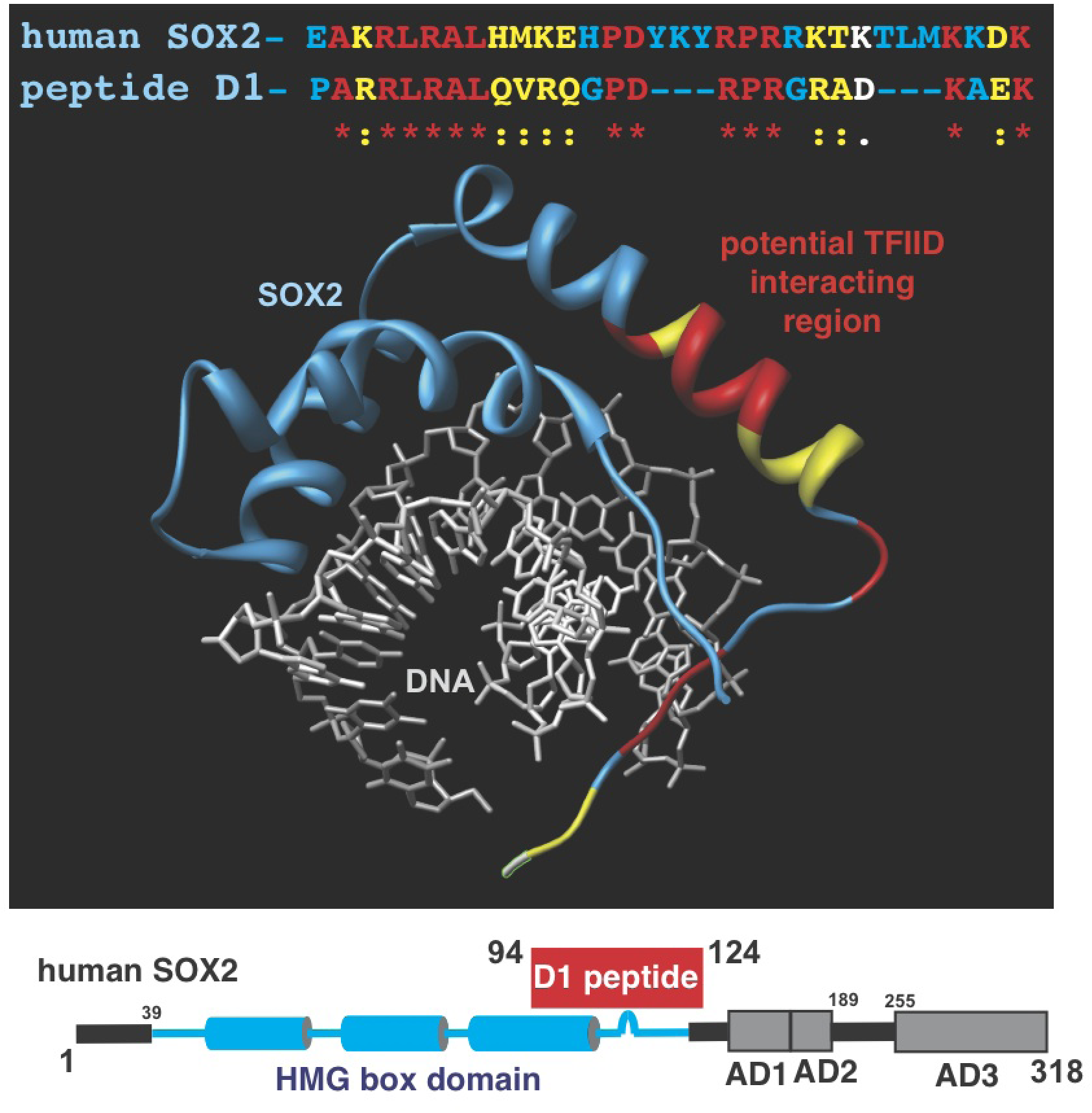
Homology between the D1 peptide and the HMG box of human Sox2. ClustalW sequence alignment of the D1 peptide and Sox 2 showing identical (red, *), highly similar (yellow, :), and similar (white, .) residues. Crystal structure (1GT0) of Sox2 (ribbon) bound to DNA (sticks) showing location of homologous residues (color coded as above) located along a-helix 3 and the c-terminal extension. Identical (red) and strongly similar (yellow) residues in peptide D1 and Sox2 are highlighted in the sequence alignment and the Sox2 (blue) X-ray crystal structure (PDB:1GT0).

**Figure 3.**
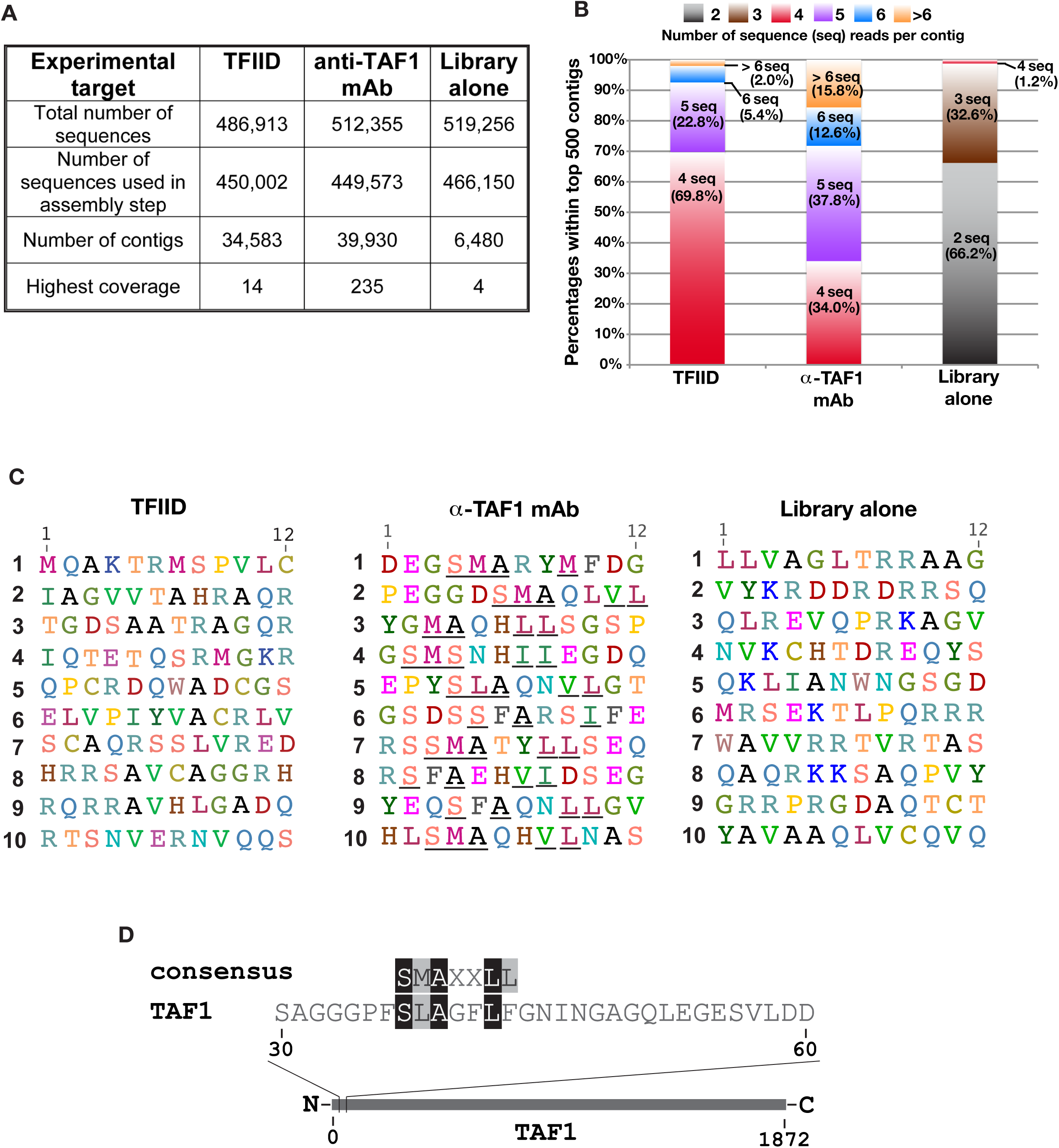
Analysis of Next Generation DNA Sequencing data from various FliTrx libraries. (A) Comparison of Next Generation sequencing data showing sequence enrichment of libraries selected with TFIID and α-TAF1 monoclonal antibodies (α-TAF1 mAb) as bait versus non-selected library alone. A contig refers to a group of identical DNA sequences. Highest coverage refers to the maximum number of identical sequencing reads within the contigs. (B) Stacked column plot showing percentage of contigs versus number of sequencing reads/contig within the top 500 contigs. The majority of contigs within the top 500 contigs contained 2 reads (66.2%), 4 reads (69.8%), and > 5 reads (66%) for the non-selected, TFIID selected and α-TAF1 mAb selected libraries respectively. (C) Peptide sequences of the top 10 contigs from the TFIID selected (left), α-TAF1 mAb selected (middle), and non-selected (right) libraries. A conserved S(L/F/M)AXXΦΦ motif (underlined) is present in all α-TAF1 mAb selected sequences. (D) Alignment of the consensus peptide sequence from the α-TAF1 mAb selected library with the amino terminus of human TAF1.

The top 10 hit peptides for each experiment are listed in Figure 3C. First of all, no overlap among these selected peptides between these different experiments was found. In addition, peptide contigs targeting the TFIID complex or the library alone did not feature any distinct patterns. These top ten TFIID- or anti-TAF1 mAb- interacting peptides could represent the most easily accessible eptiopes recognized by TFIID or the anti-TAF1 mAb due to their high sequence coverage relative to the non-selected library. Intriguingly, we noticed a specific pattern that features a characteristic S(L/F/M)AXXΦΦ motif present among the peptide contigs targeted by the anti-TAF1 mAb (middle panel). We further examined the top forty-four peptide contigs against this antibody and confirmed a concensus sequence of SMAXXLL residing within these contigs (Supplemental Figure 1). The exact epitope recognized by this anti-TAF1 antibody is unknown. However, we speculated that this selected consensus sequence could be related to TAF1. Thus, we aligned these forty-four peptide contig sequences with human TAF1 using ClustalW (Geneious, Biomatters) (Figure 3D). Remarkably, we detected that this unique residue pattern is located within the N-terminal 35∼42 amino acids of TAF1. This result suggests that the epitope of this anti-TAF1 mAb likely resides within this region.

To demonstrate the feasibility of this approach and further identify putative TFIID interactors, we next performed Blast searches against a human nuclear protein database (uniProtKB) using these top 10 peptide contig sequences. Since TFIID functions in the nucleus, we restricted our search for nuclear proteins that may associate with TFIID. The results clearly showed some known TFIID-binding factors (Table 3), such as TFIIB [3, 28] and the RPB2 subunit of RNA Pol II [6]. Interestingly, we also found some novel proteins that may target TFIID. For clarification, we sorted theses candidates into 17 classes that include Activators/Repressors, RNA Pol II, TFIIB, TFIID, TFTC, Chromatin remodelers, Histone methylase/ demethylase/ deimininase complexes, Splicing factors, Pol I transcription machinery, Ubiquitin E3 ligase, and others. It is of note that 13 overlapping and four new classes are detected with this proteomic approach compared to the previous conventional approaches (Tables 1 and 2). Importantly, there are several novel unknown TFIID interactions involved in non-transcriptional pathways such as DNA repair, kinetochore assembly and non-sense mediated decay.

**Table 3.**
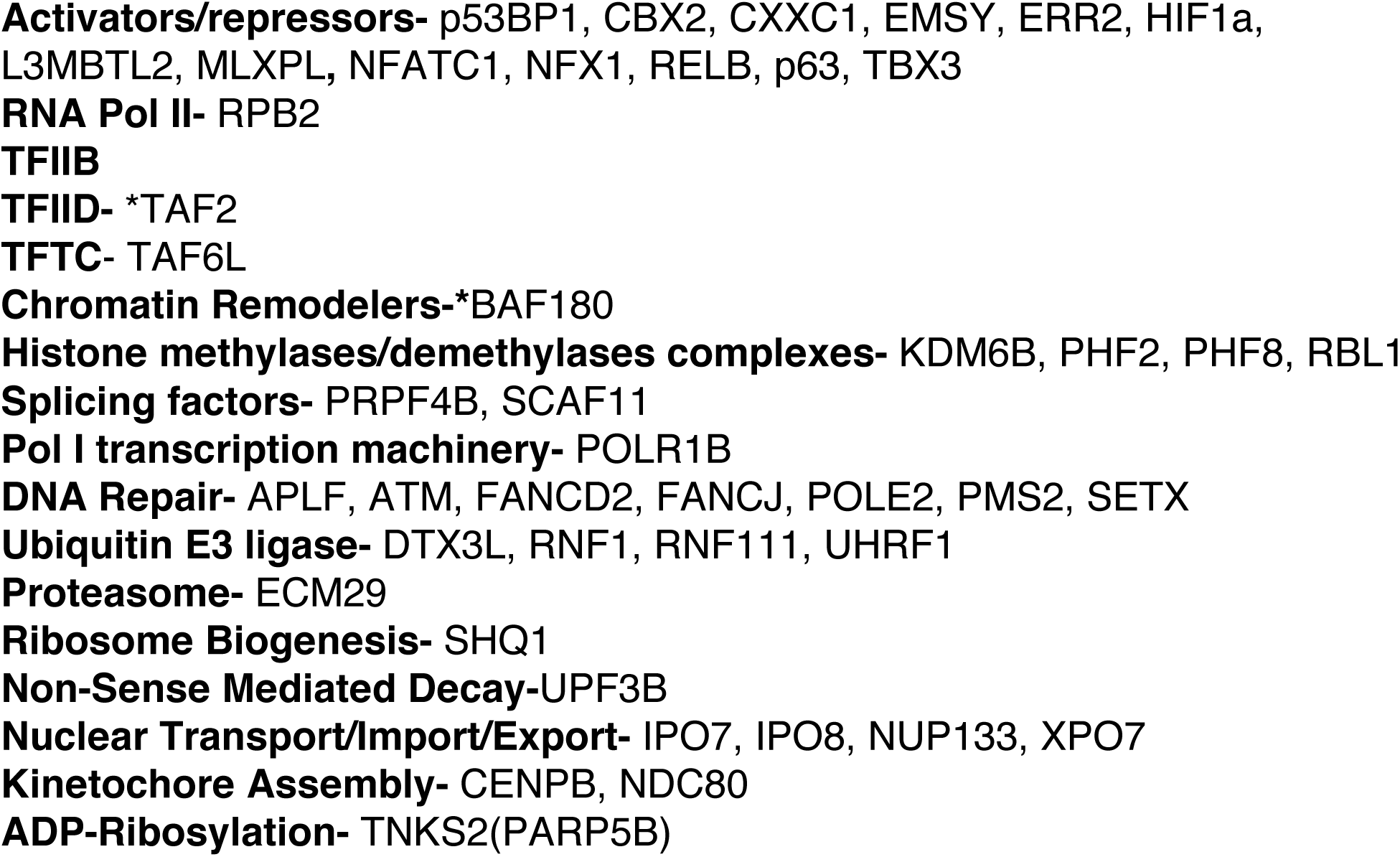
Identifying TFIID-interacting factors based on top ten peptide contigs. Blast search was conducted following the Flagella random peptide display screening and the Next generation sequencing (Ion Torrent). Based on their primary cellular functions, factors analyzed were listed in different categories. Hits also found in MudPIT are indicated with a star (*) (see Table 2 and Supplemental Table 1).

### Characterization of select TFIID-binding peptides

Early studies on the artificial acidic transcriptional activator GAL4-AH (amphipathic α-helix) demonstrated that this 15 amino acid peptide (i.e. AH) is essential to stimulate the promoter occupancy of TFIID and activate transcription [29]. In addition, multiple subunits within TFIID (i.e. TBP, TAF4, TAF6, & TAF12) cross-linked to the VP16 activation domain [30]. Hence, it would be interesting to see if our peptides can act in a similar manner to those short activating peptides that also target multiple TAFs. To assess the association between TFIID and the TFIID-interacting peptides identified from all of these approaches, we selected 12 peptides including the D1 peptide (with strong homology to a conserved region within the HMG box of Sox2) to conduct our established label transfer assays [14]. Since these 12-mer peptides are small, we tagged GST protein to these peptides for these interaction assays. These 12 peptide sequences are listed in Supplemental Table 2. With this assay we can further map potential peptide-TAF contacts in the context of intact holo-TFIID.

In brief, we performed photo-cross-linking label transfer reactions utilizing the trifunctional reagent Sulfo-SBED (S-SBED) [14]. First, we labeled each GST-peptide fusion protein or the control GST with S-SBED, respectively, using a high pH buffer to target internal tertiary amines (lysines) thus biasing the reaction towards “body” labeling. After removal of unreacted S-SBED, we incubated our highly purified TFIID with S-SBED- labeled-GST-peptide or -GST to initiate binary formation of GST or GST-peptide/TFIID assemblies. The samples were then exposed to UV to activate the aryl azide group that cross-links to nearby TAFs. After cleavage of the disulfide bond within S-SBED with DTT, the biotin moiety will be “transferred” from the GST or GST-peptide fusion proteins to any adjacent TAF within 21Å [14]. The resulting biotin-tagged TAFs in TFIID were subjected to SDS-gel electrophoresesis and examined by western blot analysis using an anti-biotin antibody.

Our analysis showed that the GST-D1 peptide most likely contacted the surface within TFIID consisting of TAF6/7 and TAF2, 3, or 4, while GST-21F predominantly binds to TAF7 but mildly cross-linked to TAF2, 3 or 4 (Figure 4A, top panel). Since TAF2/3/4 co-migrate as a single band in SDS-PAGE gels, unfortunately we couldn’t designate which specific TAF (i.e. TAF2, 3, or 4) cross-linked to the SBED-labeled GST-peptides. However, a recent study using immunopurification of Sox2 followed by MudPIT analysis revealed an interaction between Sox2 with TAF2 [31]. It is thereby likely that the GST-D1 peptide contacts TAF2 in this experiment.

**Figure 4.**
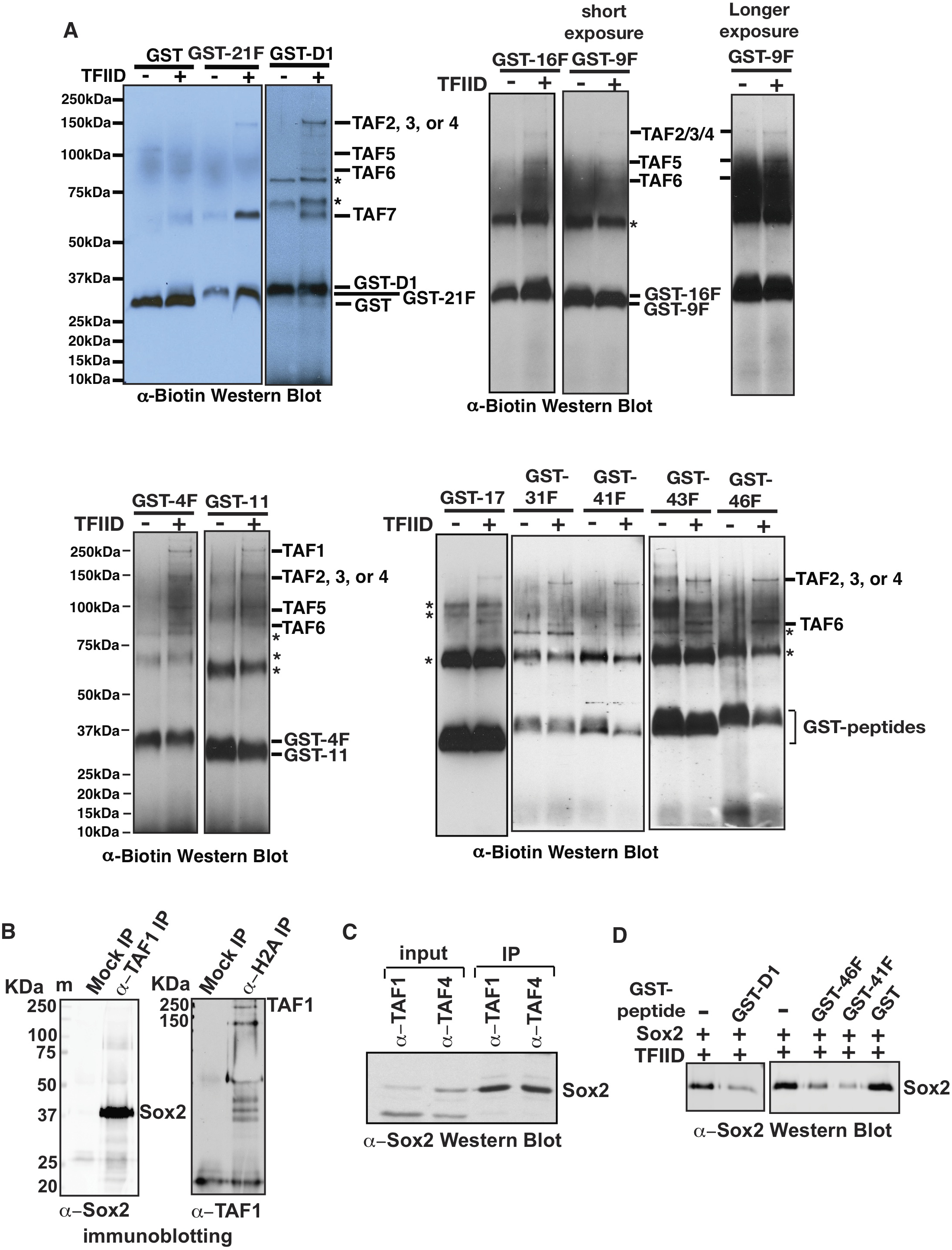
Defining TFIID’s interaction with various TFIID selected peptides and homologous full-length proteins. (A) Determination of TFIID subunits targeted by individual selected peptides using SBED label transfer assays. GST alone or GST fusions of TFIID-selected peptides were body labeled via surface exposed lysines with the SBED label transfer crosslinking reagent and incubated in the absence and presence of TFIID. Samples were exposed to UV light to initiate crosslinking and incubated with DTT to transfer biotin from the labeled GST proteins to the crosslinked TFIID subunits. SDS-PAGE followed by anti-Biotin western blotting (WB) was used to identify TFIID subunits within 21Å of the bound TFIID selected peptides. (B) Co-immunoprecipitation analysis of TFIID bound to full-length proteins containing sequences that were homologous to TFIID selected peptides. Highly purified endogenous TFIID was incubated with recombinant human Sox 2 or histone H2A. Samples were immunoprecipitated with an α-TAF1 mAb (left) or α-H2A pAb (right) followed by western blotting using antibodies against Sox2 (left) or TAF1 (right). (C) TFIID was isolated with mAbs against either the TAF1 or TAF4 subunit using nuclear extract generated from human NTERA-2 cells. TFIID precipiates were analyzed via SDS-PAGE and western blot analysis via an anti-Sox2 pAb. (D) The same co-immunoprecipitation experiments of TFIID and Sox2 were performed using α-TAF1 mAb, except that either GST, select peptide-tagged GST fusion proteins or buffer alone were incubated in each reaction as indicated in the figure. TFIID immunoprecipitates were analyzed via western blot analysis using α-Sox2 rabbit pAb. (E) TFIID was co-immunoprecipiated with EMSY from HeLa nuclear extracts using an anti-EMSY mAB.

Our label transfer data demonstrate that some peptides possibly bind similar contact surfaces within TFIID. For example, the SBED-labeled-GST-peptides 16F and 9F weakly cross-linked to bands corresponding to either TAF2, 3, or 4, in addition to TAF5/6. On the other hand, peptides 4F and 11 most likely contacted TAF1 in addition to TAF2, 3, or 4. Interestingly, we noticed that all six SBED-labeled GST-peptide fusion proteins (i.e. GST-17, -21F, -31F, -41F, -43F, and -46F) cross-linked to TAF6 along with TAF2, 3, or 4, suggesting that they may target some common surface or binding pocket within TFIID. The summary of label transfer results is listed in Supplemental Table 2. Notbaly, TFIID contains many subunits that co-fold and thereby are in close proximity within TFIID. As we used a GST-tagged 12 aa peptide (∼30kDa total) for labeling, it is possible that we surveyed a larger area (>20 Å) of TFIID with many TAFs in close proximity in our label transfer assays. Overall, our results suggest that these 12 peptides associate with native TFIID and some of these peptides may bind distinct surfaces within TFIID.

Since the D1 peptide sequence has strong homology to the Sox2 protein (Figure 2) and interacted with select TAFs (Figure 4A), we further examined if full-length Sox2 protein binds TFIID. In addition, we had noticed a strong homology between the N-terminus of histone H2A and two TFIID-interacting peptides (i.e. #1 & #3 in Figure 3C and Supplemental Figure 2) that were found to be highly enriched in our next generation sequencing analysis. Interestingly, H2A was also a positive hit present in one of the two TFIID preps using MudPIT (refer to Supplemental Table 1). Given that TFIID was previously shown to directly interact with the acetylated nucleosome tails [18], we sought to test if H2A could bind TFIID. Thus, we incubated native TFIID complexes either with purified recombinant Sox2 or H2A and then performed in vitro co-immunoprecipitation (IP) assays using antibodies against TAF1, H2A or nonspecific IgG. The co-immunoprecipitates with TFIID were visualized by immunoblotting using antibodies against Sox2 and the TAF1 subunit of TFIID as shown in Figure 4B. The results demonstrated that TFIID indeed associates with Sox2 and H2A compared to the mock IP condition.

To assess the association between TFIID and Sox2 in vivo, we immunoprecipiated TFIID from nuclear extract generated from human NTERA-2 pluripotent embryonal carcinoma using two highly specific mAbs against either the TAF1 or TAF4 subunit of TFIID [10, 14]. We also tested several non-specific IgG for the IP experiments (data not shown). TFIID precipitates were analyzed via immnuoblotting using an anti-Sox2 pAb (Figure 4C). The data showed that Sox2 associates with TFIID in these NTERA-2 cells.

Since the TFIID-interacting D1 peptide share a strong sequence homology with Sox2, we set out to test if the D1 peptide can disrupt TFIID/Sox2 interactions. Thus, we carried out the same co-IP experiment as in Figure 4B, with addition of the GST-D1 fusion protein to potentially compete TFIID/Sox2 association. In this assay, we also included two GST-peptide fusion proteins, GST-41F and GST-46F, which appeared to target somewhat similar contact surfaces within TFIID as GST-D1 (Figure 4A). As seen in Figure 4D, when we used a 5-fold molar ratio of GST-peptide:Sox2 in these reactions, a clear reduction of Sox2 interacting with TFIID was observed compared to the control GST protein. Interestingly, we found that GST-41F seemed to work better than GST-D1 in terms of competing the Sox2/TFIID association in this assay. Overall, these findings validated that the interaction between Sox2 and TFIID is specific. We further tested another novel interactor EMSY (refer to Table 3) by co-immunoprecipitaiton assays from HeLa nuclear extracts (Figure 4E). The results showed that EMSY can bind TFIID in cells. Taken together, our multi-pronged approach involving Flagella Peptide Display, Next Generation Sequencing and MudPIT mass spectrometry could permit identification of a more comprehensive mammalian TFIID interactome. This unique unbiased approach detects putative TFIID-interacting factors involved in diverse cellular pathways. We anticipate that this facile high-throughput approach can be utilized to survey the interactome of many other multi-subunit complexes.

## Discussion

A number of previous studies have used global mass spectrometry to identify factors interacting with the TBP subunit of TFIID [3, 32-34]. Alternative proteomic approaches utilizing the intact native human TFIID complex as bait to find direct interactors had not been assessed. In addition, using a bacterial peptide display screening approach to identify putative TFIID-interacting peptides/factors has not been documented. Here we exploited a random peptide library screening system on the surface of bacteria flagella to select for short peptides that interact with the surface of TFIID. In addition to a conventional DNA sequencing approach, we employed Next Generation DNA sequencing (Ion Torrent) for our analysis. We developed a data analysis pipeline using Next Generation DNA sequencing data that allows us to quantitate interactions between peptides and our target factors (i.e. TFIID and anti-TAF1 mAb). Overall, this strategy permits identification of: **i)** a more comprehensive spectrum of TFIID-binding peptides and proteins; **ii)** potential consensus TFIID-interacting epitopes; and **iii)** most common sets of TFIID-interacting factors. In this study, we only applied the top ten peptide contigs and, consequently, a limited number of candidates were found (Table 3). It is feasible to expand the search using additional top peptide contigs. Most importantly, it is critical that reasonable cutoff criteria and careful examination of the candidates must be applied when searching for novel interactors, given the fact that short 12-mer amino acid peptides are used for these analyses. Another key notion is that all of these positive candidates in this study only represent likely TFIID-interactors, which absolutely require further confirmation of their direct association with TFIID via biochemical and functional assays. Since currently a number of mass spectrometry datasets (e.g. www.crapome.org) are available, it would be interesting to further investigate those unknown putative interactors. However, when we used crapome for preliminary analysis of TFIID-interacting proteins from our MudPIT studies, we found that these hits are unique to intact TFIID immunoprecipitates and not commonly found in other mass spectrometry datasets.

Based on results obtained from both sequencing methods, we noticed some common categories of TFIID interactors, although candidates in each overlapping class are not identical (Compares Tables 1 and 3). These overlapping categories include activators/repressors, RNA Pol II, TFIID, chromatin remodelers, histone methylase/demethylase complexes, RNA Pol I transcription machinery, DNA repair factors, proteasome, ubiquitin E3 ligase, ribosome biogenesis, nuclear transport/import/export, and ADP-ribosylation (compare Tables 1 and 3). It is not surprising that most of these overlapping classes (e.g. activators/ repressors, RNA Pol II, chromatin remodelers, histone methylase/demethylase complexes) are involved in transcription regulation, considering the crucial role of TFIID in eukaryotic pre-initiation complex formation. In particular, we observed that the PBAF chromatin remodeling complex, specifically the BAF180 subunit, appeared in all of these analyses (Tables 1, 2 and 3). This result is consistent with the previous report showing that the PBAF complex tightly co-purified with TFIID through multiple column purification steps [35]. In contrast, yeast RSC, a homolog of the human PBAF complex, solely interacts with TBP and not TFIID as evidenced by the lack of RSC subunits in immunoprecipitates of TAF subunits [3]. This suggests that humans may have evolved to utilize PBAF to remodel nucleosomes present at a wider variety of promoters (i.e. TFIID regulated) versus yeast (i.e. TBP/SAGA regulated).

The prime interest for future studies will be to examine if TFIID indeed associates with factors participating in other cellular processes such as DNA repair, proteasome, and ubiquitin E3 ligases. Thus far, MDM2 is the only ubiquitin E3 ligase that has been shown to directly interact with mammalian TFIID via the TAF1 and TBP subunits [36, 37]. A previous study on native yeast TFIID revealed co-purification of the ubiquitin E3 ligase Rsp5 with the observation of TAF1 and TAF5 being ubiquinated [34]. In addition, multiple human TFIID subunits (TAF1, 2, 7, 9 and TBP) are ubiquitinated by unknown E3 ligases [38, 39]. Thus, it is likely that human TFIID could interact with one or multiple of these E3 ubiquitin ligases found in our analysis.

### Interactions between TFIID and the RNA Pol III transcription machinery

Both MudPIT and Flagella peptide screening/sequencing analyses have come up with human TFIID-interacting candidates involved in RNA Pol III transcription. It is possible that interactions between TFIID and RNA Pol III factors may regulate either RNA Pol II or Pol III directed transcription. Interestingly, previous ChIP-seq studies revealed that additional TFIID interacting factors such as, TFIIA, TFIIB, TFIIE and RNA polymerase II are found at Pol III regulated promoters in vivo [40]. TFIIIB, TFIIIC, and RNA Pol III can also be found at or adjacent to Pol II regulated promoters [40-43]. Furthermore, it is possible for Pol III to transcribe these traditional Pol II regulated genes [40]. However, the role of TFIID in regulation of transcription by RNA Pol III remains an enigma. Given that the TBP subunit is required for both Pol II and Pol III transcription, it is possible that a surface on TBP utilized by Pol III factors is accessible in TFIID. Yet TBP may not be the only TFIID subunit targeting RNA Pol III factors. Immunoprecipitations of yeast TAF5 also yielded RNA Pol III subunits suggesting that interactions between TFIID and the Pol III machinery may be conserved [3].

### Similarities between the human and yeast TFIID interactomes

A number of reports documented that different TAFs within TFIID bind to different transcriptional activators to regulate select subsets of gene expression programs [6]. However, TFIID is also known to bind transcriptional repressors such as N-Cor and RBPJ (CBF1) via TAF6/9 or TAF4 respectively [44, 45]. This suggests that TFIID utilizes a variety of surfaces (i.e. TAF subunits) to interact with gene specific regulators that may be either transcriptional activators or repressors. In addition to targeting various activators, it is possible that TFIID may contact other factors responsible for regulating different cellular processes related to gene expression. A previous comprehensive proteomics study on yeast TFIID has revealed that different TAFs and TBP bind a varied number of factors involved in different cellular pathways including RNA processing, signal transduction, and RNA Pol III transcription [3]. Interestingly, we have identified many identical targets as found in the yeast TFIID proteomics study [3]. For example, we also find that TFIID interacts with human MED14, TFIIB (i.e. yeast SUA7), BAF180 (i.e. yeast RSC1, 2, and 4), RPL23, H2A, and HIRA in either our MudPIT or flagella display assays. Conservation of these TFIID associated factors in both of our studies suggest potential physiologically relevant interactions with TFIID.

### Correlation between TFIID subunits targeted by selected peptides and known TFIID interactors

Our initial SBED label transfer studies were able to localize specific peptide interactions with select TAFs (Figure 4A). For example, our D1 peptide crosslinked primarily to TAF7 along with TAF2, 3, or 4 (Figure 4A). This D1 peptide has strong homology to both Sox2 (Figure 2) and TIF1A (Supplementary Figure 3A), which are transcriptional co-regulators [46, 47]. Intriguingly, a recent study documented an interaction between Sox2 with TAF2 via immunopurification of Sox2 followed by MudPIT analysis [31], which is consistent with our D1-peptide label transfer results (Figure 4A). We have also confirmed a direct interaction between Sox2 and TFIID using co-immunoprecipitation studies (Figures 4B & 4C). In vitro assays show that TIF1A can phosphorylate TAF7 [22], another TAF that was crosslinked to the D1 peptide (Figure 4A), suggesting that our label transfer assays may measure physiologically relevant interactions.

Peptide 21F primarily crosslinks to TAF7 (Figure 4A). Blast searches of peptide 21F reveal homology with several factors involved in DNA repair including DNA Ligase I, a human endonuclease III like factor (NTHL1), and Cyclin D1(CCND1) [48] (Supplemental Figure 3B). TAF7 was previously found to interact with TFIIH, a factor that is also involved in DNA repair along with DNA Ligase I [49, 50]. TAF7 is also known to dissociate from TFIID upon transcription initiation and may travel along with RNA Polymerase II via its ability to interact with the elongation factor pTEF-b [49]. Thus we speculate that when RNA Pol II encounters a region of damaged DNA, TAF7 may be a conduit for recruitment of DNA repair factors to repair the DNA damage.

We have noticed that many of our peptides found through conventional (Figure 4A) and Next Generation sequencing (data not shown) were crosslinked to a band corresponding to either TAF2, 3, or 4 on SDS-PAGE gels of TFIID. Due to the limit of the label transfer assays, we couldn’t define which TAF among these three TAFs serves as a common site for interaction with many regulators of TFIID function. However, many transcriptional regulators such as activators and repressors bind to TAF4 [14, 51]. TAF4 contains an ETO-TAFH domain that is known to interact with multiple activators and repressors [51]. In addition, TAF4 contains a histone fold domain that resembles histone H2A [52], which could also be targeted by transcriptional regulators. Thus, it is likely that TAF4 contains major regulatory within TFIID.

Overall, based on the variety of potential TFIID interactors, this work could provoke future studies in co-regulatory pathways linked to transcription initiation. In addition to identifying novel TFIID-interacting factors, common TFIID-interacting peptide sequences can allow us to delineate regions within these factors that may be physiologically important. This localization of potential interaction surfaces in our TFIID-interacting factors will be the starting point that allows the facile generation of mutants to impair these interactions in vivo. Thus we anticipate using this information to determine the in vivo function of these interactions. In addition, this high-throughput sequencing proteomic approach can be easily adapted for other proteins of interest without certain constraints such as transient protein-protein interactions, cell type-specific expression and low abundance of interactors.

## Materials and Methods

### Purification of TFIID

Human HeLa cells (32 liters) were grown in suspension culture with 1X DMEM plus 5% newborn calf serum. Nuclear extract was prepared as previously described and fractionated with phosphocellulose P11(P-Cell) resins [10]. P-Cell column fractions eluting at 1M KCl/HEMG buffer (pH 7.9, [20 mM Hepes, 0.2 mM EDTA, 2 mM MgCl_2_, 10% glycerol] plus 1 mM DTT and 0.5 mM PMSF) were pooled, dialyzed to 0.3 MKCl/HEMG buffer (plus 0.1% NP40 and 10 μM leupeptin) and immunoprecipitated overnight at 4°C with an anti-TAF4 monoclonal antibody (mAb) covalently conjugated to protein G sepharose beads (GE Healthcare Life Sciences). TAF4-immunoprecipitates were extensively washed with 0.65 M KCl/HEMG buffer, 0.3 M KCl/HEMG buffer and 0.1M KCl/HEMG buffer (containing 0.1% NP40 buffer and 10 μM leupeptin). The TFIID complexes from our TAF4 mAb affinity resin were eluted with a peptide (1 mg/ml in 0.1 M KCl/HEMG buffer/0.1% NP40) recognized by the TAF4 mAb. The eluates were concentrated with a microcon-10 concentrator.

### Flagella peptide display screen

The FliTrx Random peptide display screening system contains 2x10^8^ clones of a random 12-residue peptide library displayed on the bacterial cell surface Flagella (Invitrogen). The experiments were conducted following manufacturer’s instruction except with minor modifications. In brief, 750 ng of highly purified native human TFIID complex was placed on a tissue culture dish prior to incubation of the primary FilTrx peptide library bacterial culture. After incubation, we gently washed away unbound bacteria and then collected the remaining bound cells by vortexing. We then grew up the eluted bacteria and used this pooled culture to re-incubate with surface attached TFIID followed by collecting the TFIID-bound cells. To enrich the peptide pool specifically bound to TFIID, we performed another four rounds of panning as described above. Single colonies were obtained from the fifth panning. In order to reveal the TFIID-interacting peptides, plasmid DNA was either extracted from each colony and subjected to conventional DNA sequencing, or from the mixed pooled TFIID-enriched bacteria followed by massive parallel Next Generation sequencing (Ion Torrent).

Specific for Next Generation sequencing, mixed pool plasmid DNA for each experiment (i.e. TFIID, anti-TAF1 mAb, and library alone) was labeled with different barcodes individually with a PCR amplication following manufacturer’s instructions. The primers used for the reactions are (1). 5’-CCTCTCTATGGGCAGTCGGTGATGGGCGATCATTTTGCACGGACC-3’ (2). 5’-CCATCTCATCCCTGCGTGTCTCCGACTCAGAGCACTGTAGATTTCTGGGCAGAGTG GTGC-3’ (for TFIID) (3). 5’-CCATCTCATCCCTGCGTGTCTCCGACTCAGACGGTCGAC-AATTTCTGGGCAGAGTGGTGC-3’ (for anti-TAF1 mAb) (4). 5’-CCATCTCATCCCTGCGT-GTCTCCGACTCAGACGAGTGCGTATTTCTGGGCAGAGTGGTGC-3’ (for library). The PCR amplication was performed with 1 cycle of 98°C for 2 min, 25 cycles of 50°C for 30 seconds, 72°C for 15 seconds, 98°C for 30 seconds, and the final cycle of 72°C for 10 mins. PCR reactions were purified with 1:1 ratio volume of phenol/CHCl_3_ extraction and subject to precipitation with 70% ethanol. The precipitated DNA was analyzed with 1.8% agarose electrophoresis. The 147 bp PCR fragment for each experiment was purified using illustra GFX PCR DNA and Gel band purification kit (GE healthcare) following manufacturer’s instructions. Libraries were amplified on beads in an emulsion PCR using the Ion Xpress Template Kit and protocol. One hundred base sequencing was performed using a 316 chip on the Ion PGM machine per protocol provided with the kit (Life Technologies, Carlsbad, CA) at the Genomics Core Facility of the Huck Institutes of the Life Sciences at Penn State University.

### MudPIT analysis

The MudPIT analysis was conducted previously and the detailed protocol was described in [10]. To identify strong-hit peptides, we set the filtered conditions with presence in two repeat immunoprecipitations, Normalized Spectral Abundance Factors (NSAF) values above 10^-4^, and 5 times more than control NSAF.

### Peptide sequence data analysis

Data obtained from the Next Generation sequencing were catalogued with Geneious Pro 5.5.6 software (Biomatters). Readouts were initially separated by the barcodes introduced during PCR amplification. Sequences shorter than 70 bp were removed from the dataset and all remaining sequences were trimmed to a maximum of 100 bp from the 5’ end. Sequences were then assembled using Geneious Assembly tool with Medium Sensitivity/Fast settings (15% gaps per read with 2 bp maximum gap size allowed, word length 14 bp, index word length 12 bp, words repeated more than 200 times ignored, reanalyze threshold 8, maximum mismatched per read 15%, maximum ambiguity 4, pair read distances used to improve assembly). Assembled contigs, built by the largest number of sequences, were then translated (genetic code 15) and manually inspected for the peptide sequences that displayed a correct pattern of dodecamer random library peptide in the context of upstream and downstream sequences (CGPXXXXXXXXXXXXGPC). Peptide hits targeting the TFIID complex did not show any clear pattern. On the other hand, upon inspection of the peptide sequence data for selection with anti-TAF1 mAb, peptides featuring a characteristic S(L/F/M)AXXΦΦ sequence (present in 27 out of 30 top assemblies) were identified. These peptide hits were then aligned with each other along with the human TAF1 protein sequence using ClustalW built into Geneious software with standard conditions (cost matrix: BLOSUM, gap open cost 10, gap extend cost 0.1). The top 10 peptide hits were presented to the PSI-BLAST program (Position-Specific Iterated BLAST) using a database (uniProtKB) consisting of human nuclear proteins to identify potential protein candidates these peptides might represent. Top 10 hits with the largest sequence coverage that represent interesting and likely interacting partners were chosen for further studies.

### Purification of peptide-tagged GST fusion proteins

Since these 12-mer peptides are small, we extended the peptide sequences by keeping the N-terminal two extra residues from the vector, inserting one cysteine residue at the N-terminus and two lysine resides at the C-terminus of the peptide sequences, and then fused with a GST tag for label transfer assays and peptide competition/immunoprecipitation analysis (see below). pGEX6p-peptide was transformed into BL21 DE3 cells and incubated at 37°C for overnight. A single colony was selected and inoculated into 3 ml of LB medium with 200 μg/ml of Ampicillin followed by shaking the culture at 220 rpm 37°C for overnight. After this step, 150 μl was taken from this culture to 150 ml of LB medium with 200 μg/ml of Ampicillin and incubated as previously. 25 ml of this pre-culture was transferred into 500 ml of TB medium and the bacteria concentration at O.D. 600nm was measured. The culture was grown until the O.D. reached 0.6∼0.8. To induce the expression of these GST-peptide fusion proteins, IPTG was added to a final concentration of 1 mM. The culture continued to grow for 3 hours at 30°C. Cells were collected by centrifugation at 5,000 rpm for 10’ at 4°C. The cell pellets were washed once with LB medium and once with ice-cold 1XPBS. After the wash, cells were lysed in 20ml of ice-cold 1XPBS containing 100 mg of lysozyme. The lysates were nutated at 4°C for 10’ and then frozen in liquid nitrogen. The frozen pellets were thaw and sonicated for 2 min on ice. The soluble fraction was collected after centrifugation at 14K rpm for 25’ at 4°C. This supernatant containing overexpressed GST-tagged peptide fusion proteins was incubated with 100 μl of Glutathione-Sepharose 4B agarose resins (GE Healthcare) at 4°C for 4 hours. After spinning down the resins, the beads were washed five times with ice-cold 1MKCl/1xPBS plus 0.2 mM PMSF. Next, the resins were washed eight times with ice-cold 0.5M KCl/1xPBS plus 0.2 mM PMSF, followed by two washes with ice-cold 1xPBS plus 0.2 mM PMSF. GST-tagged peptide proteins were eluted off the resins with 100 μl of elution buffer (10 mM reduced L-Glutathione [Sigma] in 100 mM KCl/1xPBS buffer plus 0.1% NP-40 and 20 mM Tris-HCl [pH 7.9]) at 4°C for 15’. GST-tagged peptide eluates were analyzed with 15% SDS-PAGE and Colloidal Coomassie Blue staining.

### Label transfer assays

1 μg of GST alone or GST-tagged peptide in Buffer L (pH 7.9 [20 mM Hepes, 100 mM KCl, 10% Glycerol, 2 mM MgCl_2_, 10 μM TCEP, 0.2 mM PMSF, 0.1% NP-40]) along with a mock control (0 μg of GST/GST-tagged peptide) were incubated with 15 μl of glutathione agarose resin (GE Healthcare) for 1 hour at 4°C and extensively washed to remove unbound proteins. The trifunctional Sulfo-SBED crosslinker (Thermo Scientific, 0.5 mM in 100 mM KCl/HEMG buffer [pH 7.9, 20 mM Hepes, 0.2 mM EDTA, 2 mM MgCl_2_, 10% glycerol]) was added to react with lysines on the resin bound GST/GST-tagged peptides or mock control at room temperature in the dark for 15’. Unreacted crosslinker was removed by extensive washing with 100 mM KCl/HEMG buffer. SBED-conjugates were eluted off the resins with 40 μl of elution buffer (10 mM reduced L-Glutathione [Sigma] in 138 mM KCl/HEMG buffer in the dark at 4°C for 20’.

80 ng of Sulfo-SBED labeled GST/GST-tagged peptide proteins were incubated with 1μl of TFIID (conc. ∼200 ng/μl) at 30°C for 45 min in the dark. The mixture was then exposed to UV light (365 nm) at room temperature for 10 min to activate the aryl azide moiety on the SBED labeled GST or GST-peptide proteins for covalent crosslinking of any protein within 21Å of the GST/GST-tagged peptides thus transferring the biotin moiety from peptides to the crosslinked subunits of TFIID. The reactions were treated with DTT to cleave the disulphide bond. Crosslinked TAFs were resolved on 4-12% Bis-Tris NuPAGE gels (Thermo Fisher) and subjected to immunoblotting using anti-Biotin antibody (Rockland) using 1:3000 dilution. The identity of biotin labeled TAFs was determined based on their known migration in 4-12% Bis-Tris Gels and confirmed by silver staining. Each reaction was repeated at least three times with representative data shown.

### In vitro co-immunoprecipitation and TFIID-interacting peptide-tagged GST competition assays

125 ng of human native TFIID complex was incubated with 500 ng of purified human Sox2 (Stemgent) on ice for 1 hour. For assessing the interaction between H2A and TFIID, 635 ng of human native TFIID complex and 500ng of purified human Sox2 were used. After this step, 1 μg of non-specific IgG Ab, anti-TAF1 mAb or anti-H2A rabbit polyclonal Ab (pAb, Active Motif) was added to each reaction, respectively. Reactions were incubated on ice for 1.5 hours. Next, 20 μl of protein G beads (GE Healthcare) resuspended in 70 μl of 150 mM KCl/HEMG buffer (pH 7.9, [20 mM Hepes, 2 mM MgCl_2_, 0.2 mMEDTA, 10% glycerol, 0.1%NP-40, 1mM DTT, 0.2mM PMSF]) were added into each mixture and nutate at 4°C for 1 hour. The resins were washed three times with 150 mM KCl/HEMG buffer. Immunoprecipitates were resolved on either 12% or 15% SDS-PAGE electrophoresis and subjected to immnuoblotting analysis using anti-Sox2 rabbit pAb (1:1000 dilution, Thermo Scientific Pierce) or anti-TAF1 rabbit pAb (1:1000). Each reaction was repeated at least three times with representative data shown.

183 ng of TFIID and 500 ng of purified human Sox2 (Stemgent) were incubated with 2.5 μg of GST-tagged peptide proteins at 4°C for 1.5 hours. Protein samples were immuno-precipitated with anti-TAF4 mAb-conjugated protein G beads at 4°C for 2.5 hours to pull down TFIID and its associated factors in each reaction. Resins were washed three times with 150 mM KCl/HEMG buffer to remove the unbound proteins. TFIID-precipitates were analyzed with 12 % SDS-PAGE electrophoresis and immnuoblotting using anti-Sox2 rabbit pAb (1:1000, Thermo Scientific Pierce). Each reaction was repeated at least four times with representative data shown.

### In vivo immunoprecipitation

Human NTERA-2 pluripotent embryonal carcinoma cells were grown in 1X DMEM plus 10% FBS. Thirty 15cm^2^ dishes of cells were harvested for further purification. Nuclear extract was prepared as previously described [10, 53]. 3.6 mg of nuclear extracts for each IP reaction was pre-cleared with 9 μl of protein G resins (GE Healthcare) at 4°C for 40’. The pre-cleared nuclear extract was incubated with either anti-TAF4 or anti-V5 mAb (Life Technoglies) conjugated protein G beads and nutated at 4°C for 5 hours. Resins were washed extensively with 100 mM KCl/HEMG buffer (pH 7.9, [20 mM Hepes, 2 mM MgCl_2_, 0.2 mMEDTA, 10% glycerol, 0.05%NP-40, 1mM DTT, 0.2mM PMSF, 10 μM Leupeptin]). The precipitates were resolved in 12% SDS-PAGE and subjected to Western Blot analysis using ant-Sox2 pAb (Cell Signaling).

## Acknowledgement

We specifically thank Robert Tjian for his support as the initial work was performed in his laboratory. In addition, we thank M. Washburn and L. Florens for the MudPIT analysis. We also thank S. Zheng for providing TAF4 mAb supernatant, D. King for peptides, and L. Woodley for the initial label transfer work. We appreciate C. Kenworthy for critical comments of this manuscript. This work was supported by the startup funds from Albert Einstein College of Medicine (W.L. and R.C.).

## Author Contributions

R.C. and W.L. designed experiments, analyzed data and wrote the paper. L. S. conducted experiments. G. D. generated reagents and performed experiments. A. P. analyzed data.

## Nomenclature, abbreviations, units and symbols

HMG: High Mobility Group
mAb: monoclonal antibody
MudPIT: Multidimensional Protein Identification Technology
NSAF: Normalized Spectral Abundance Factors
pAb: polyclonal antibody
TAF: TBP-associated factor
TBP: TATA-binding protein
TFIID: Transcription factor IID

